# Neural correlates of individual variability in reward-based motor learning

**DOI:** 10.1101/2025.08.06.668596

**Authors:** Hayato Otake, Naoki Senta, Junichi Ushiba, Mitsuaki Takemi

## Abstract

Individual differences in motor learning are linked to variability in the neural processes associated with feedback processing and motor memory retention. This study investigated the neural correlates of reward-based motor learning and focused on two electrophysiological markers: event-related desynchronization (ERD) in the sensorimotor cortex and feedback-related negativity (FRN) from frontal EEG channels. Sixty-four healthy adults performed a visuomotor rotation task with monetary reward or punishment feedback. Learning amount, defined as the degree of error compensation, and retention amount, defined as the maintenance of adapted motor behavior in a no-vision condition, were quantified. Scalp EEG was recorded to assess alpha (8–13 Hz) and beta (15–30 Hz) ERD over the primary motor cortex (M1) during movement preparation and FRN amplitude following feedback presentation. Stepwise regression revealed that FRN amplitude in the late adaptation phase was associated with learning amount, but only among participants with robust event-related potential (ERP) responses (amplitude > ±5 µV). This finding suggests that greater neural sensitivity to feedback supports better learning. In contrast, alpha ERD magnitude in the late adaptation phase, along with reinforcement conditions, was a significant predictor of motor skill retention in the full sample, although no interaction effect was observed. These findings suggest that FRN and alpha ERD reflect dissociable neural mechanisms underlying learning and retention, respectively, with FRN indexing feedback sensitivity and alpha ERD reflecting cortical excitability related to memory consolidation. The absence of interaction effects between neural markers and reinforcement conditions further supports the idea that these processes contribute independently to motor adaptation. Practically, combining reward-based training with interventions that increase M1 excitability may enhance retention, while tailoring feedback strategies to maximize FRN responses may improve learning efficiency. These findings provide novel insights into the neural basis of individual variability in motor learning and suggest avenues for personalized approaches in rehabilitation and skill learning.

## 1. Introduction

Rewards influence learning processes by shaping both procedural and declarative memory formation (Chen et al., 2018; Miendlarzewska et al., 2016). They can be both positive and negative, and they activate the reward circuitry in the brain through the dopamine system (Zhao et al., 2024a). Comparing positive rewards and punishments (i.e., negative rewards), positive rewards increase the likelihood of repeating a behavior by reinforcing it and motivating continued effort, while punishments decrease the likelihood of a behavior by promoting avoidance of adverse outcomes (Schultz et al., 1997). These differing motivational effects have been demonstrated in motor learning contexts, where rewards have been shown to reduce forgetting and enhance retention, whereas punishments can increase the speed of adaptation (Galea et al., 2015).

Importantly, individual variability is commonly observed in the extent to which rewards and punishments affect behavior, likely due to underlying neural differences. For example, inter-individual differences in reward-based learning have been linked to baseline levels of striatal dopamine, with higher dopamine levels enhancing reward-based learning and lower levels enhancing punishment-based learning (Cools et al., 2009). Likewise, variability in neural responses to the same amount of monetary rewards has been reported during reinforcement learning tasks (Kim et al., 2015). Such variability in brain activity may also shape the extent of behavior modification in motor learning, including the speed of learning and retention of motor memory (Zhao et al., 2024a).

Given the potential for rewards to enhance the effectiveness of interventions in motor rehabilitation and athletic training (Grau-Sánchez et al., 2020; Quattrocchi et al., 2017; Zhao et al., 2024b), it is important to understand the neural factors that drive individual differences in motor learning efficiency. To this end, we recorded scalp electroencephalogram (EEG) data while participants performed a visuomotor rotation task in which monetary rewards or punishments varied according to the size of their motor error (Galea et al., 2015). While EEG offers millisecond temporal resolution for tracking neural dynamics during motor preparation, execution, and feedback phases, its limited spatial resolution poses challenges for localizing the underlying sources. To address this, we focused on two EEG features whose cortical origins and functional interpretations are well established: sensorimotor event-related desynchronization (ERD) in the alpha and beta bands, which reflects primary motor cortex (M1) excitability (Takemi et al., 2013; Takemi et al., 2018), and frontal feedback-related negativity (FRN), an event-related potential (ERP) linked to feedback processing and predominantly generated in the anterior cingulate cortex (ACC) (Gehring & Willoughby, 2002; Holroyd & Coles, 2002; Miltner et al., 1997). These EEG features provide non-invasive proxies for monitoring cortical processes relevant to motor execution and feedback evaluation.

In this study, we examined how M1 and ACC activities contribute to individual variability in motor learning and retention. Specifically, we assessed M1 excitability using sensorimotor ERD during movement preparation and evaluated feedback processing using frontal FRN amplitude following reward or punishment feedback. Learning amount, defined as the degree of error compensation during adaptation, and retention amount, defined as the persistence of compensation without visual feedback, were separately quantified. By examining these electrophysiological markers across different learning phases, we aimed to identify whether cortical excitability and feedback processing independently account for inter-individual differences in motor learning performance.

## 2. Materials and Methods

### 2.1. Participants

Sixty-four young adults (24 ± 5 years, 12 females) participated in this study. The target sample size was determined a priori using G*Power 3.1 (Faul et al., 2009). Effect-size estimates were derived from two relevant EEG studies. Palidis et al. (2019) reported *R*² = 0.242 for a multiple-regression model linking FRN components to motor learning performance, whereas Pollok et al. (2014) found a correlation of *r* = -0.67 (*r*² ≈ 0.45) between beta band ERD changes and reaction-time improvement. We assumed that there were modest inter-predictor correlations among the seven planned regressors (reward condition plus six EEG variables) and conservatively set the expected overall coefficient of determination to *R*² = 0.25 (equivalent to *f*² = 0.33). A fixed-model multiple-regression power analysis (*α* = 0.05, 1-*β* = 0.80, 7 predictors) indicated a required sample of 51 participants. We anticipated a 20% data loss based on a pilot experiment (5% attrition and 15% unusable EEG recordings), so we recruited 64 participants to preserve the desired statistical power.

All participants were right-handed, had no history of neurological or psychiatric diseases, and were not on chronic medication. Five participants who responded on the questionnaire that they were unable to perform the task properly due to drowsiness or lack of motivation were excluded from the results. Data from seven participants were excluded from the EEG analysis due to poor signal quality caused by body movements or other noise (*n* = 4) or malfunctions in the EEG recording system (*n* = 3), although their data were included in the behavioral analysis. The study was conducted in accordance with the Declaration of Helsinki, and the research protocol was approved by the ethics committee of the Faculty of Science and Technology at Keio University (IRB approval numbers: 2023-071 and 2024-010). All participants provided written informed consent.

### 2.2. Experimental apparatus

Scalp EEG signals were recorded using an EEG System (GES 400; Electrical Geodesics, Inc., Oregon, USA) equipped with a 128-channel HydroCel Geodesic Sensor Net (HCGSN-128) at a sampling rate of 1,000 Hz. The ground and reference electrodes were positioned at CPz and Cz, respectively, following the extended 10-20 system. Electrode impedance was maintained below 50 kΩ. The EEG signals were filtered online using a band-pass filter (0.1 to 70 Hz) and a notch filter (50 Hz).

A custom-designed program developed in C++ was used to implement the motor task. Participants sat with their chin resting on a chin rest and grasped the handle of a robotic manipulandum (Phantom Premium 1.5HF, SensAble Technologies, Wilmington, MA) with their right hand in a semi-pronated forearm posture (Fig. 1A). A computer monitor (1920 × 1080 pixels) positioned above the manipulandum appeared to be in the same plane as the participant’s hand, which effectively obscured direct visual feedback of the hand position. Instead, a cursor representing the hand’s position was displayed on the monitor, with its positional information calculated based on the manipulandum’s movements. Handle position data were recorded at a sampling rate of 1,000 Hz.

**Figure 1.**
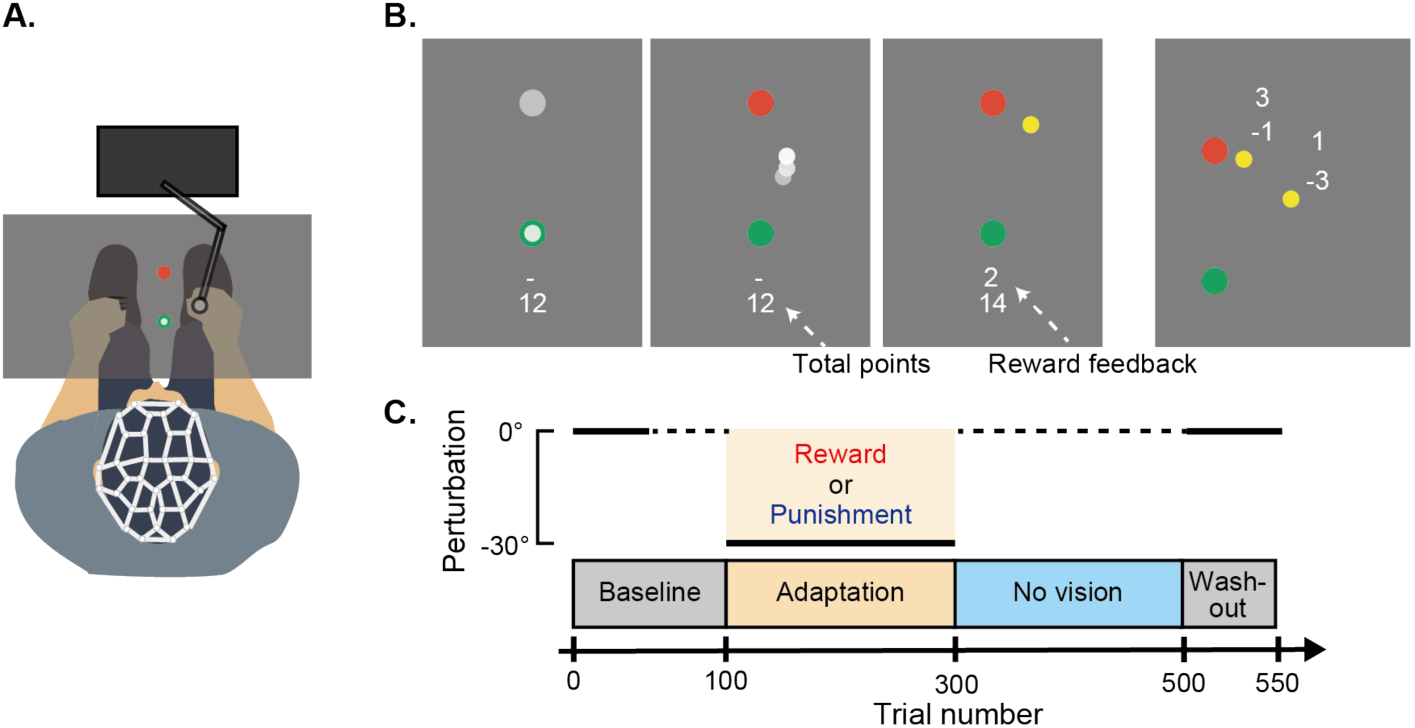
Experimental design. (A) Experimental setup. Participants performed arm-reaching movements toward a visual target displayed on a monitor while wearing an EEG cap to record brain activity. (B) Experimental task. Reaching movements started from a green circle (starting position) to a red circle (target). During each trial, participants received online feedback of their hand position (white circle) as well as endpoint feedback (yellow circle). Reward or punishment feedback was represented by positive or negative points based on endpoint error, with points accumulated throughout the adaptation block and displayed on the monitor. (C) Experimental paradigm. The task consisted of four blocks: baseline, adaptation, no vision, and washout. Online and endpoint feedback were provided for the baseline (first 50 trials), adaptation, and washout blocks. During the adaptation block, a 30° counterclockwise visuomotor rotation was introduced, and reward or punishment feedback was provided.

### 2.3. Motor task

The motor task required participants to manipulate a cursor displayed on the monitor using a robotic manipulandum to perform shooting movements toward a visual target (Fig. 1B). At the beginning of each trial, participants moved a white cursor (0.3 cm diameter) into a green circle (1 cm diameter) representing the starting position. Once the cursor was held within the starting position for more than 1.5 s, the monitor displayed a gray target (0.5 cm diameter) located 8 cm in front of the starting position. After a fixed interval of 1 s, the target changed color from gray to red, serving as the Go cue. Participants were instructed to make a fast and accurate shooting movement through the target to minimize online corrections during the movement. When the movement stopped, the location where the cursor crossed the invisible boundary of an 8-cm radius circle centered on the starting position was displayed with a yellow circle to indicate the endpoint error. Additionally, feedback regarding movement speed was provided. Movements with maximum speeds below 312 mm/s were labeled as “Slow,” while those exceeding 469 mm/s were labeled as “Fast.” After each trial, participants moved the cursor back to the starting position (Galea et al., 2015).

The experiment consisted of four blocks: baseline, adaptation, no vision, and washout (Fig. 1C), with visual feedback varying across blocks. During the baseline block (trials 1 to 100), the cursor was visible, and endpoint feedback was provided for the first 50 trials; however, both cursor and endpoint feedback were removed in the latter 50 trials. In the adaptation block (trials 101 to 300), a 30° counterclockwise visuomotor rotation was applied, causing the cursor’s movement to deviate from the hand’s movement. This visuomotor transformation introduced a motor error, which required participants to modify their hand trajectories to compensate for the altered environment. In the no-vision block (trials 301 to 500), participants performed reaching movements without any visual feedback. During this phase, no information about movement accuracy was provided, allowing hand trajectories to gradually return to baseline, thereby characterizing the degree of memory retention. Finally, the washout block (trials 501 to 550) replicated the conditions of the first 50 trials of the baseline block.

To prevent fatigue, participants were given short rest periods of less than 3 min every 100 trials. During these breaks, they were instructed to keep their arms beneath the monitor. EEG signals were recorded continuously throughout all experimental blocks.

### 2.4. Visuomotor rotation with reward and punishment feedback

Participants were divided into two groups: the reward group (*n* = 32) and the punishment group (*n* = 32). During the adaptation block, the type of feedback varied depending on the group, with the reward group receiving positive feedback and the punishment group receiving negative feedback (Table 1). The reward group started with 0 points, while the punishment group started with 800 points. Both groups could earn a maximum of 800 points and a minimum of 0 points, with points accumulating throughout the block. The reward group accumulated positive points, while the punishment group accumulated negative points. Participants in both groups were shown the points they earned on a trial-by-trial basis, along with the total accumulated points for the block. The total points accumulated by the end of the block were converted into Japanese yen at a rate of three yen per point, and the resulting amount was added to their participation wage. All participants were informed that closer reaches to the target would result in a greater monetary incentive, and their final score would determine their additional compensation. They were also made aware of the maximum and minimum points they could earn, as well as the conversion rate of points to yen. Participants received a base wage of 1,000 yen/h, to which their bonus earnings were added.

**Table 1.**
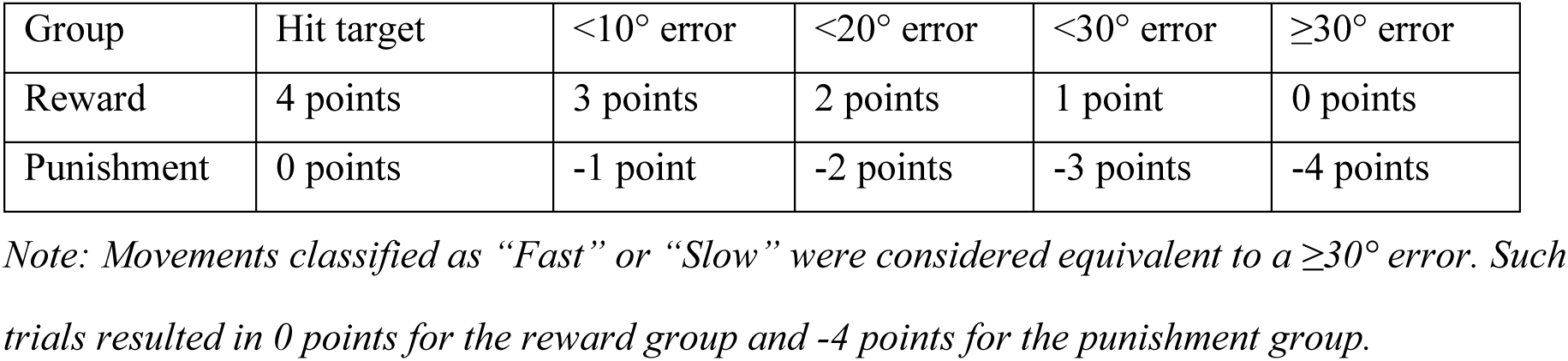
Point system for reward and punishment conditions.

### 2.5. Behavioral data analysis

Behavioral data analysis was conducted using MATLAB R2023a (MathWorks, Natick, MA, USA). For each trial, the angular reach direction was calculated as the angular difference between hand and target positions at the point where the cursor crossed the 8-cm invisible circle centered on the starting position. Under veridical feedback conditions, the ideal reach direction was 0°. However, during the visuomotor transformation phase, participants were required to compensate for the rotation. For example, with a -30° counterclockwise visuomotor rotation, accurate compensation required a 30° clockwise reach direction.

To quantify motor learning and retention, we defined two metrics: the learning amount, which reflects the degree of motor memory acquisition, and the retention amount, which reflects the degree of motor memory retention. These metrics were calculated using the following equations:

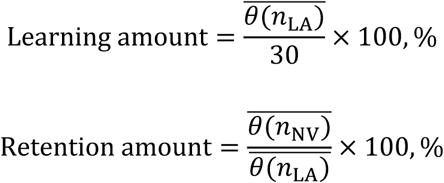

Here, *θ*(*n*) represents the reaching direction at trial *n*, *n*_LA_ refers to trials in the late adaptation phase (trials 201 to 300), and *n*_NV_ refers to trials in the no-vision phase (trials 301 to 500).

### 2.6. EEG preprocessing

EEG analysis was performed using MATLAB R2023a and the FieldTrip toolbox (Oostenveld et al., 2011). Raw EEG signals were filtered offline using a 4th-order Butterworth filter, with different frequency ranges applied depending on the analysis: 0.1 to 30 Hz for FRN analysis and 1 to 60 Hz for ERD analysis. Noisy channels were excluded based on their power spectral density and amplitude range. A common average reference method was applied as a spatial filter, wherein the average signal across all EEG channels was subtracted from each channel to reduce noise and improve signal quality. Independent component analysis was then conducted to remove artifacts caused by eye movements and muscle activity. Artifact-related independent components were manually identified and subtracted from the EEG signals. Finally, spatial interpolation was conducted using the FieldTrip toolbox “ft_channelrepair” function to reconstruct signals for the excluded channels. This function uses weighted data from neighboring channels.

### 2.7. Event-related desynchronization analysis

Cleaned EEG data were segmented into 7 s epochs, time-locked to the Go cue (-3 to +4 s). Trials containing visual noise or artifacts were manually identified and excluded. The power spectral density (PSD) was calculated using the FieldTrip toolbox “ft_freqanalysis” function with the multitaper convolution (mtmconvol) method. For PSD computation, a 1 s Hanning-tapered sliding window was used, advancing in 0.1-s steps (90% overlap) to provide high temporal resolution for ERD analysis. ERD was computed using the following equation:

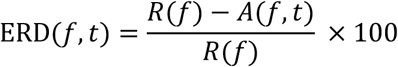

Here, *R*(*f*) represents the average PSD during the reference period (from 2 to 1 s before the Go cue) at a frequency *f*, and A(*t*, *f*) denotes the PSD at a given time point *t*.

ERD values were calculated for each participant by averaging data over the left primary motor cortex (channel 36) and its six surrounding channels. The maximum ERD values were identified within the time range of -1 to 0 s relative to the Go cue, corresponding to the preparatory phase of the reaching movement. These maximum ERD values were analyzed separately for the alpha band (8 to 13 Hz) and beta band (15 to 30 Hz). Average ERD values were then computed within a ±0.2 s time window and ±2 Hz frequency range centered on the maximum ERD values. The resulting averaged ERD values were subsequently used for statistical analyses.

### 2.8. Feedback-related negativity analysis

FRN is most prominently observed in EEG signals recorded from FCz (Holroyd & Krigolson, 2007; Miltner et al., 1997; Pfabigan et al., 2011) and is commonly analyzed using the difference-wave approach in ERPs. FRN is classically characterized by a negative voltage deflection following non-reward feedback. However, recent studies suggest that reward-related positive responses, while not forming a distinct peak alone, also contribute to FRN when analyzed using the difference-wave approach,. This analysis allows for a more accurate representation (Baker & Holroyd, 2011; Becker et al., 2014; Carlson et al., 2011; Heydari & Holroyd, 2016; Walsh & Anderson, 2012).

In this study, cleaned EEG signals recorded from FCz and its surrounding electrodes (channels 5, 6, 7, 12, 13, 106, and 112), corresponding to the area above the ACC, were segmented into 0.8 s epochs, time-locked to the onset of the score feedback stimulus (-200 to +600 ms). Trials containing visual noise or artifacts were manually identified and excluded. ERPs were calculated by separately averaging the EEG time series across trials for successful outcomes (reward group: 4 points; punishment group: 0 points) and unsuccessful outcomes (reward group: 3 to 0 points; punishment group: -1 to -4 points). Baseline correction was applied to all ERPs by subtracting the average voltage during the 75 ms period immediately following the onset of the score feedback from each trial. This post-feedback baseline period was chosen because it corresponds to a time when participants were not actively performing motor tasks, which resulted in relatively stable EEG voltages. Additionally, the latency of this period is too short to be influenced by visual feedback processing, making it an optimal period for baseline correction. After applying baseline correction to individual trials, ERPs were averaged across all trials within each condition (e.g., successful and unsuccessful outcomes) to calculate condition-specific waveforms. For each participant, difference waves were calculated by subtracting ERPs for unsuccessful outcomes from those for successful outcomes. The FRN amplitude was then determined as the mean voltage of the difference waves within the 200 to 350 ms time window following the presentation of score feedback, in line with a previously established protocol (Palidis et al., 2019).

### 2.9. Statistical analysis

Statistical analyses were conducted using JASP (version 0.18.3; https://jasp-stats.org/). To compare reach direction, ERD values, and FRN amplitudes between reward and punishment feedback conditions and across experimental blocks, a two-way ANOVA was conducted with block as the within-subject factor and feedback condition as the between-subject factor. Prior to conducting ANOVA, we assessed the sphericity assumption using Mauchly’s test. If the sphericity assumption was violated, we applied the Greenhouse–Geisser correction to adjust for the degrees of freedom. If ANOVA yielded statistically significant main effects or feedback × block interactions, post-hoc t-tests were performed with Bonferroni correction to compare specific condition or block differences. A p-value of less than 0.05 was considered statistically significant.

## 3. Results

### 3.1 Effects of reward and punishment on visuomotor rotation task performance

Figure 2A illustrates the results of the visuomotor rotation task. The adaptation and no-vision phases were further divided into early and late phases, and the mean angular reach directions were analyzed for these four phases (Fig. 2B). A two-way repeated-measures ANOVA revealed a significant interaction effect between phase (early adaptation to late no vision) and feedback condition (reward vs. punishment). Mauchly’s test of sphericity indicated a violation of the sphericity assumption (*p* < 0.05). Consequently, the Greenhouse–Geisser correction was applied, confirming a significant interaction effect between phase and feedback condition (*F*(1.3, 75.0) = 7.31, *p* = 0.005). Post-hoc pairwise comparisons identified significant differences between the reward and punishment groups in both early and late phases of the no-vision block (*p* < 0.001). Substantial inter-subject variability was observed in the no-vision phase, as reflected by the shaded areas in Fig. 2A and the large error bars in Fig. 2B. An independent samples t-test revealed no significant difference in total scores between the reward and punishment groups (*t*(57) = 0.70, *p* = 0.49), indicating that overall task performance during the adaptation phase was comparable between the two feedback conditions (Fig. 2C).

**Figure 2.**
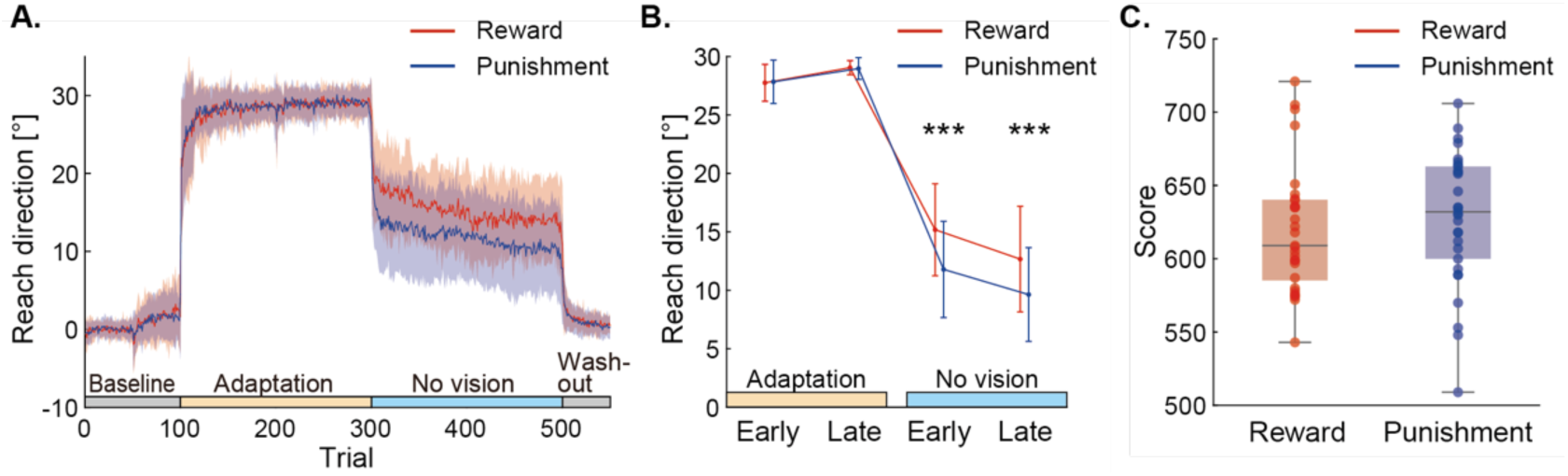
Effects of reward and punishment on motor adaptation and retention. (A) Trial-by-trial angular reach direction during the visuomotor rotation task for the reward group (*n* = 29) and punishment group (*n* = 30). Solid lines indicate mean values, while shaded areas represent standard deviations. (B) Mean reach direction during the early and late phases of the adaptation and no-vision blocks. Significant differences between the reward and punishment groups were observed in both the early and late no-vision phases (\*\*\**p* < 0.001). Error bars indicate standard deviations. (C) Total scores for the reward and punishment groups. Box plots display the median, interquartile range, and individual data points.

### 3.2 ERD changes during the adaptation task

Figure 3A shows the time-frequency representation of ERD during the adaptation task, focusing on the alpha and beta frequency bands. ERD during the preparatory phase (-1 to 0 s relative to task onset) was observed across nearly all participants, irrespective of the reward or punishment condition (Fig. 3B). Alpha and beta ERD during the early and late adaptation phases were compared between the reward and punishment groups. For alpha ERD, a two-way repeated-measures ANOVA revealed a significant main effect of time (*F*(1, 50) = 4.90, *p* = 0.031), indicating a change in ERD magnitude between the early and late adaptation phases. However, there was no significant interaction between time and reward condition (*F*(1, 50) = 2.02, *p* = 0.16), nor was there a significant main effect of reward condition (*F*(1, 50) = 0.12, *p* = 0.73). Similarly, for beta ERD, a two-way repeated-measures ANOVA revealed a significant main effect of time (*F*(1, 50) = 4.92, *p* = 0.031), suggesting a time-dependent change in ERD magnitude. However, no significant interaction between time and reward condition was observed (*F*(1, 50) = 0.014, *p* = 0.91), and the main effect of reward condition was also not significant (*F*(1, 50) = 0.98, *p* = 0.33). These results indicate that the observed ERD changes during the adaptation task were primarily driven by temporal factors rather than by the type of feedback condition.

**Figure 3.**
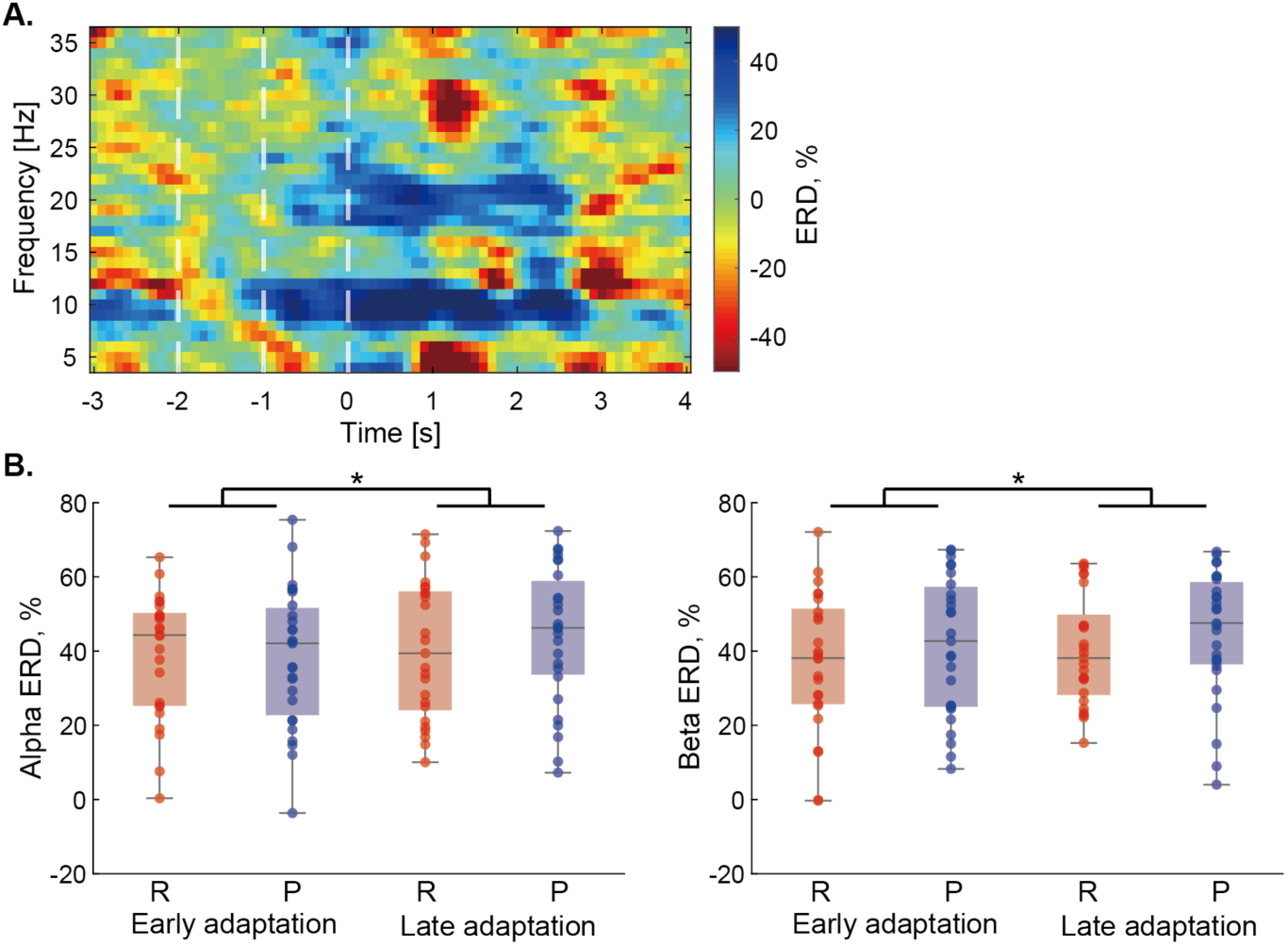
Event-related desynchronization (ERD) during motor adaptation is not influenced by reward or punishment. (A) Time-frequency map of a representative participant during the adaptation task, measured over the left primary motor cortex. The preparatory phase (-1 to 0 s) is highlighted by white dotted lines. ERD was calculated relative to the baseline power spectrum (-2 to -1 s). (B) Alpha (8–13 Hz) and beta (15–30 Hz) ERD magnitudes during the preparatory phase in the early and late adaptation phases for both the reward group (R, *n* = 25) and punishment group (P, *n* = 27). Box plots display the median, interquartile range, and individual data points. Significant main effects of time were found for both alpha and beta ERD (*: *p* < 0.05), suggesting a time-dependent change in ERD magnitude.

### 3.3 FRN changes during the adaptation task

During the adaptation task, ERPs associated with FRN varied across participants. A subset of participants (reward group: *n* = 10, punishment group: *n* = 6) exhibited ERP amplitudes exceeding ±5 μV in response to score feedback (Fig. 4A), while the remaining participants showed smaller or negligible ERP responses. Given that ERPs should differ significantly from background EEG activity to be considered a genuine neural response (Picton et al., 2000), we restricted statistical comparisons between the reward and punishment groups to participants with ERPs exceeding ±5 μV (Fig. 4B). A two-way repeated-measures ANOVA revealed a significant main effect of time (*F*(1, 14) = 6.74, *p* = 0.021), indicating that FRN amplitudes changed between the early and late adaptation phases. However, no significant interaction was observed between time and reward condition (*F*(1, 14) = 1.47, *p* = 0.25), and no significant main effect of reward condition was detected (*F*(1, 14) = 0.013, *p* = 0.91). These findings suggest that changes in FRN amplitudes were influenced by temporal factors rather than the type of feedback condition.

**Figure 4.**
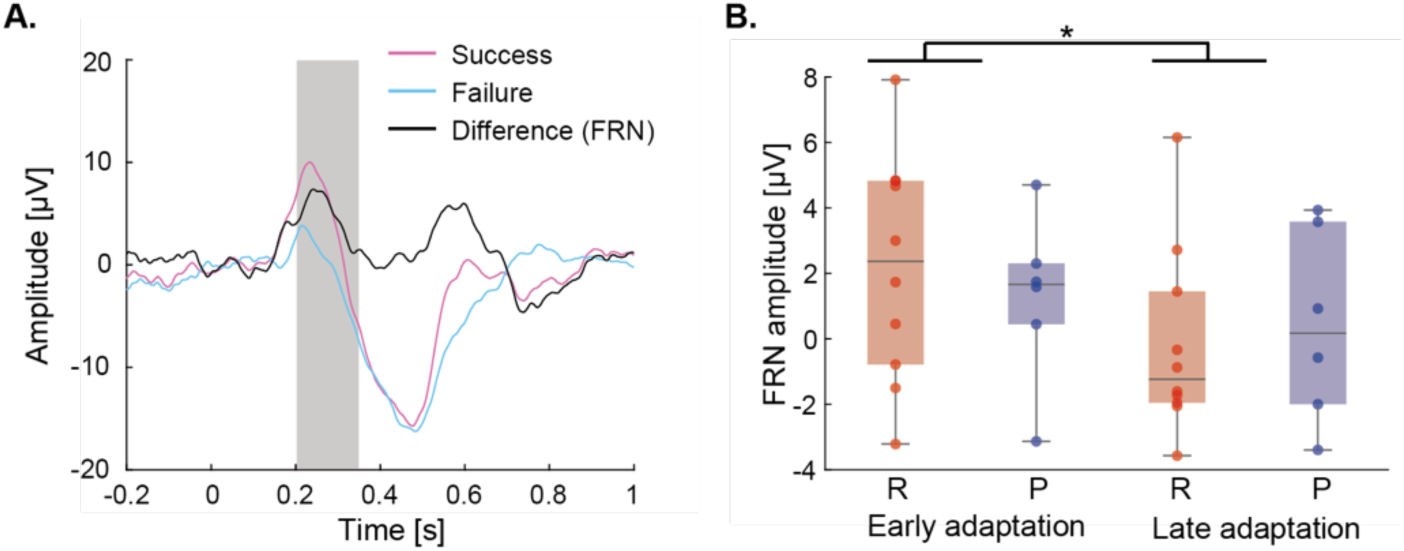
Feedback-related negativity (FRN) during adaptation tasks is not influenced by reward or punishment conditions. (A) Event-related potentials (ERPs) during the early adaptation task for one representative participant. The black waveform represents FRN, calculated as the difference between ERPs for success and failure trials. Time zero marks the onset of feedback presentation (reward or punishment). (B) FRN amplitudes during the early and late adaptation phases for participants with pronounced ERP responses (greater than ±5 μV). Box plots display the median, interquartile range, and individual data points of the reward group (R, *n* = 10) and punishment group (P, *n* = 6). A significant main effect of time was found (*: *p* < 0.05), suggesting a time-dependent change in FRN amplitudes.

To address concerns about potential selection bias due to the exclusion of participants with low-amplitude ERPs, we conducted an additional analysis using data from all participants whose EEG recordings were deemed valid (*n* = 52). Statistical comparisons of FRN between the reward and punishment groups for all participants are shown in Supplementary Fig. 1A. A two-way repeated-measures ANOVA again revealed a significant main effect of time (*F*(1,50) = 4.08, *p* < 0.05), indicating changes in FRN amplitudes over time. However, no significant interaction was found between time and reward condition (*F*(1,50) = 2.73, *p* = 0.11), and no significant main effect of reward condition was detected (*F*(1,50) = 0.30, *p* = 0.59).

### 3.4 Neural correlates of learning and retention in motor adaptation

To identify neural correlates of individual differences in learning and retention, we conducted stepwise regression analyses using MATLAB R2023a. The following variables were included as regressors: reward-punishment condition (categorical), FRN amplitudes during early and late adaptation phases, and alpha and beta ERD magnitudes during early and late adaptation (Fig. 5A).

**Figure 5.**
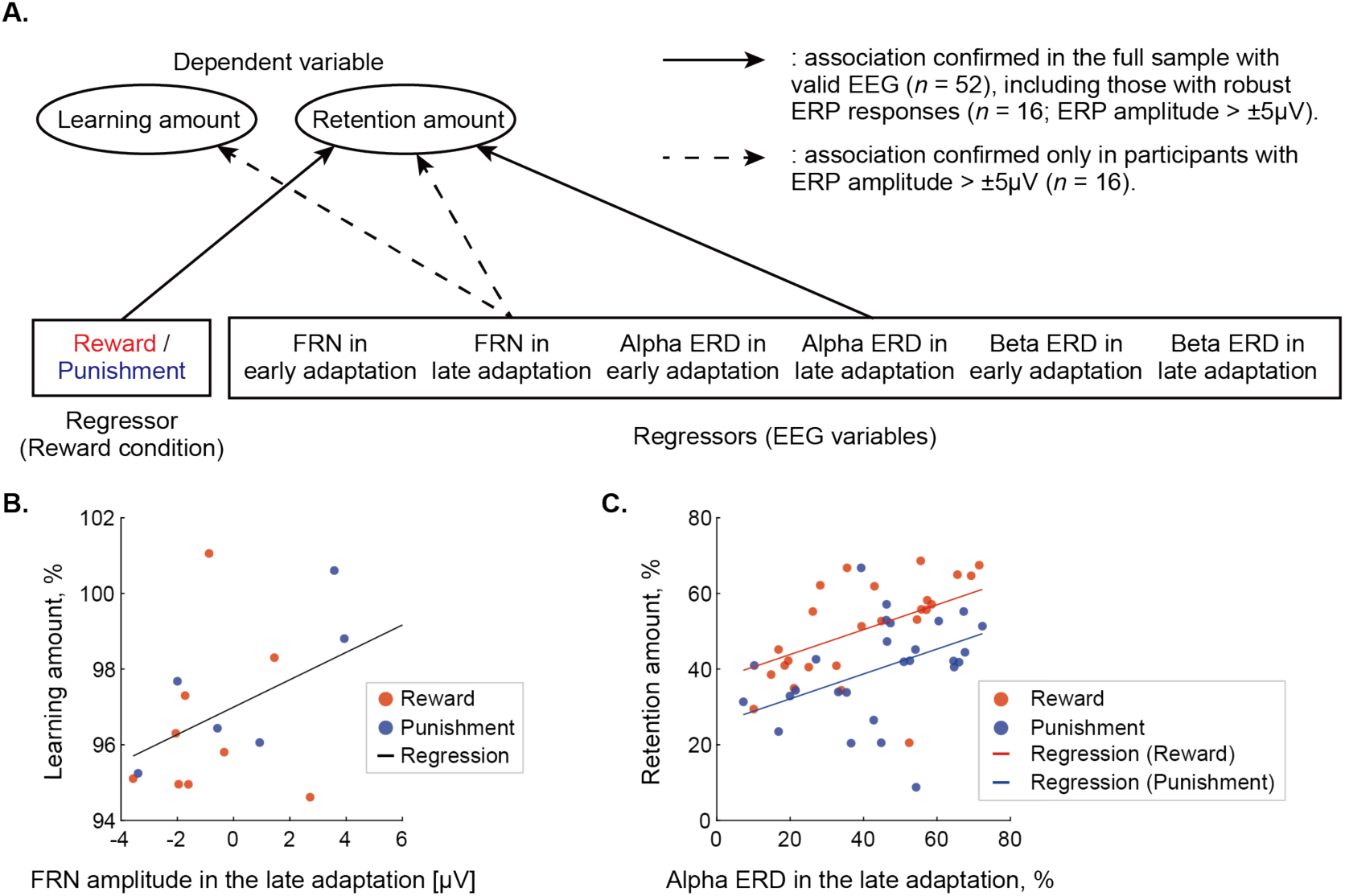
Neural correlates of learning and retention in motor adaptation. (A) Schematic of the regression models. Arrows indicate significant associations identified through stepwise regression. Solid arrows represent associations observed in the full sample (*n* = 52), while dashed arrows indicate associations confirmed only in participants with robust ERP responses (*n* = 16; ERP amplitude > ±5 μV). (B) Scatter plot showing the relationship between FRN amplitude during the late adaptation phase and learning amount. Data are from participants with ERP amplitudes > ±5 μV (*n* = 16). (C) Scatter plot showing the relationship between alpha ERD in the late adaptation phase and retention amount, separated by reward condition. Regression lines represent model fits for each group. Data are from all participants with valid EEG (*n* = 52).

For the learning amount, two separate regression analyses were conducted. The first used data from all participants with valid EEG recordings (*n* = 52), including those with robust ERP responses (*n* = 16; ERP amplitude > ±5 µV) and those with weaker responses (*n* = 36; ERP amplitude ≤ ±5 µV). This analysis revealed no significant predictors of learning amount, suggesting that FRN and ERD measures do not explain individual differences in learning when the full sample is included.

The second analysis was restricted to participants with robust ERP responses (*n* = 16). In this subset, the best-fitting model identified FRN amplitude during the late adaptation phase as a significant predictor (Adjusted *R*^2^ = 0.196, *p* = 0.049; Fig. 5B):

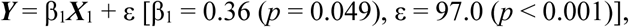

where ***Y*** denotes learning amount, ***X***_1_ is FRN amplitude in the late adaptation phase, and ε is the error term. This result highlights that larger FRN amplitudes were associated with higher learning amounts.

A similar approach was applied to identify predictors of retention. In the first model using data from all participants (*n* = 52), alpha ERD magnitude in the late adaptation phase and reward-punishment condition emerged as significant predictors (Adjusted *R*² = 0.300, *p* < 0.001; Fig. 5C):

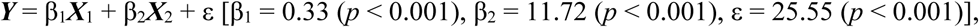

where ***Y*** represents retention amount, ***X***_1_ is alpha ERD magnitude in the late adaptation phase, ***X***_2_ is reward-punishment condition, and ε is the error term. No interaction between ERD and reward-punishment condition was observed, suggesting that cortical excitability (indexed by ERD) and reward-based reinforcement signals act as independent, additive contributors to retention.

The second regression model, using only the subset with robust ERP responses (*n* = 16), identified alpha ERD, FRN amplitude, and reward condition as significant predictors of retention (Adjusted *R*² = 0.862, *p* < 0.001):

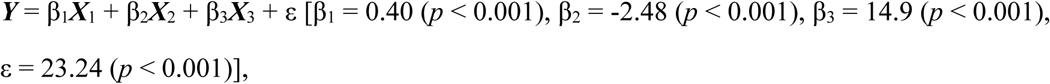

where ***X***_1_ is alpha ERD magnitude in the late adaptation phase, ***X***_2_ is FRN amplitude in the late adaptation phase, and ***X***_3_ is reward-punishment condition.

## 4. Discussion

This study investigated the neural correlates of individual differences in reward-based motor learning, with a focus on ERD over sensorimotor regions and FRN measured from frontal EEG channels. Stepwise regression analyses revealed that FRN amplitude during the late adaptation phase significantly predicted learning amount, but only in participants who exhibited robust ERP responses. This finding suggests that individuals with heightened cortical sensitivity to score feedback and greater neural differentiation between success and failure exhibited superior learning performance. Additionally, both reward-punishment condition and alpha ERD magnitude in the late adaptation phase were significant predictors of motor memory retention across all participants, with no interaction between these factors. These results imply that long-term retention may be optimized by combining positive rewards with heightened M1 excitability during motor learning. Overall, this study highlights FRN and alpha ERD as potential neural markers of individual variability in motor learning and retention. This supports the notion that distinct neural mechanisms underlie learning acquisition and memory consolidation.

### 4.1. Alpha ERD in the late adaptation phase and its association with motor memory retention

Our results revealed a clear association between alpha ERD in the late adaptation phase and motor memory retention. However, the precise functional role of alpha ERD in motor memory remains an open question. Below, we discuss several possible mechanisms that may explain why greater alpha ERD is linked to improved motor skill retention.

One possible explanation is that alpha ERD reflects increased excitability in M1 (Takemi et al., 2013), which may facilitate the long-term retention of learned motor skills. This hypothesis aligns with previous findings, which showed that enhancing M1 excitability through anodal transcranial direct current stimulation during motor learning improves motor memory retention (Galea et al., 2011; Reis et al., 2009). Another potential mechanism involves the suppression of task-irrelevant information and the optimization of task-relevant neural activity. Alpha-band oscillations have been implicated in attentional filtering, selectively inhibiting distracting information while enhancing neural processing relevant to the task (Klimesch et al., 2007; Jensen & Mazaheri, 2010). This attentional modulation may contribute to the stabilization of motor memory, thereby improving skill retention.

Alternatively, stronger alpha ERD in the late adaptation phase may not cause, but rather reflect, improved motor memory retention. Individuals who develop more refined sensorimotor representations may exhibit greater ERD, which indicates increased activation of cortical neuronal networks (Neuper & Pfurtscheller, 2001), which in turn enhances motor memory retention. The significant increase in alpha and beta ERD from the early to the late adaptation phase observed in our study aligns with this interpretation. Prior research also indicates that M1 excitability, as indexed by ERD, increases with continued motor practice (Gehringer et al., 2019; Nelson et al., 2017), possibly reflecting a shift from cognitively demanding control to more automatic execution. In the early adaptation phase, ERD tends to be modest as neural activity is primarily driven by error monitoring and rapid feedback-based adjustments (Holroyd & Coles, 2002). Over time, as learning progresses, motor control becomes more internally regulated, potentially giving rise to stronger ERD signals (Gehringer et al., 2019; Nelson et al., 2017). This shift may reflect a consolidation process in which sensorimotor representations become more robust, facilitating long-term retention.

Although our findings established an association between alpha ERD and motor memory retention, the causal relationship remains unclear. An important next step is to determine whether direct modulation of alpha oscillations can enhance retention. This could be tested using non-invasive brain stimulation techniques such as rhythmic transcranial magnetic stimulation (Thut et al., 2011) or transcranial alternating current stimulation (Wach et al., 2013), allowing for a causal assessment of alpha-band activity in motor memory consolidation.

### 4.2. FRN and individual differences in motor learning

Stepwise regression analysis identified FRN amplitude in the late adaptation phase as a significant predictor of learning amount, but only among participants who exhibited robust ERP responses. This finding suggests that individuals with stronger cortical sensitivity to score feedback and greater neural differentiation between successful and unsuccessful outcomes may demonstrate superior motor learning. FRN is thought to reflect reward prediction errors, guiding adaptive behavior through reinforcement learning mechanisms (Holroyd & Coles, 2002). The observed relationship between FRN and learning amount is consistent with this framework. This indicates that individuals who exhibit heightened neural responsiveness to performance feedback may be better at refining motor strategies.

However, this association was not observed when the regression analysis was conducted across all participants, including those with weaker ERP signals. Furthermore, no significant difference in learning amount was found between participants with larger ERP responses (> ±5 μV; *n* = 16) and those with smaller responses (*n* = 36) (*t*(50) = 0.45, *p* = 0.36; Supplementary Fig. 1B). These results suggest that alternative learning mechanisms, such as sensory-based error correction, may compensate for weaker ERP responses. Participants who relied more on score feedback likely exhibited larger ERP amplitudes, whereas those prioritizing sensory prediction errors may have shown weaker responses. Future studies should incorporate subjective reward value ratings to assess how perceived feedback relevance influences neural and behavioral outcomes.

The observed decline in FRN amplitude over the adaptation phase suggests a gradual shift in learning strategies. As motor execution efficiency improves, participants may become less dependent on external feedback (Holroyd & Coles, 2002). However, this reduction in FRN amplitude could also result from habituation rather than learning-related improvements. Given that FRN is most pronounced when errors are unexpected (Heldmann et al., 2008), a progressive decrease in error salience may contribute to its attenuation over time. Distinguishing between these possibilities remains a challenge and warrants further investigation.

### 4.3. Limitations of the study

A key limitation of this study is the absence of punishment-induced acceleration in learning, which has been reported in previous research (Galea et al., 2015). One possible explanation is that, although participants received negative score feedback, many may have relied more on sensory error feedback, which would reduce the influence of punishment. This view is supported by the finding that only a small subset of participants exhibited a robust FRN response. This suggests that punishment feedback may not have been a primary driver of learning for most individuals. Additionally, how punishment was framed may have affected its effectiveness. Unlike previous studies where punishment involved explicit monetary loss, our study used score-based feedback, which may have been perceived as less consequential. A clarification of how variations in punishment framing influence learning dynamics would help refine reinforcement-based training protocols.

Another limitation is the spatial resolution of EEG, which restricts its ability to assess activity in deep brain structures such as the striatum, a key region in reward-based learning. While EEG provides valuable insights into cortical activity, it cannot directly capture neural processes within subcortical structures that may contribute to individual differences in learning and retention. Future studies could use functional MRI or transcranial focused ultrasound stimulation to complement EEG findings and investigate how reward processing in deeper brain regions influences motor learning outcomes (Nakajima et al., 2022).

Lastly, caution is warranted when interpreting the negative association observed between FRN amplitude and retention amount in the subset of participants with robust ERP. While the FRN amplitude was positively associated with learning amount, it was negatively associated with retention. However, this inverse relationship is likely an artifact of the retention calculation. As a result, larger learning values can mathematically yield smaller retention scores, even when actual performance remains consistent. Thus, the observed negative association between FRN and retention may not reflect a true neural mechanism but rather the structure of the dependent variable.

## 5. Conclusions

This study identified distinct neural markers associated with individual differences in motor learning outcomes. Specifically, alpha ERD in the late adaptation phase was a robust predictor of motor memory retention, while FRN amplitude in the same phase predicted learning amount, though the latter association was observed only in participants with pronounced ERP responses. These findings suggest that motor learning and retention involve partially dissociable neurophysiological mechanisms. Alpha ERD likely reflects processes related to motor memory consolidation through enhanced M1 excitability, whereas FRN reflects sensitivity to performance feedback during the learning phase.

Importantly, our results indicate that individual variability in motor skill acquisition and retention may arise from intrinsic neurophysiological traits, such as cortical excitability (indexed by ERD) and responsiveness to feedback (indexed by FRN), rather than from external reinforcement conditions alone. The absence of significant interaction between ERD, FRN, and reinforcement conditions supports the idea that these neural processes contribute independently rather than synergistically. Additionally, the progressive increase in alpha and beta ERD, along with the systematic decrease in FRN amplitude over time, suggests a gradual transition from reliance on external feedback to more internally driven motor control. This pattern aligns with established theories of motor learning and consolidation (Schmidt, 1975; Wulf & Lewthwaite, 2016).

From a practical perspective, optimizing motor learning and retention may involve integrating reward-based training with interventions that enhance M1 excitability, such as non-invasive brain stimulation. This approach holds promise for applications in rehabilitation and athletic training. Additionally, feedback strategies that maximize FRN responses at the individual level could improve motor learning efficiency and offer a potential framework for personalized training interventions.

## Acknowledgments

The authors thank Dr. Seitaro Iwama (Faculty of Science and Technology, Keio University) for his valuable comments on this work.

## Data availability statement

Data used to generate all plots in the figures can be found at https://osf.io/u5wvn/. Other data (e.g., raw data) will be provided upon request to the corresponding author.

## Author contributions

Hayato Otake: formal analysis, investigation, software, visualization, writing – original draft Naoki Senta: methodology, software, validation, visualization, writing – review & editing Junichi Ushiba: funding acquisition, resources, supervision, writing – review & editing Mitsuaki Takemi: conceptualization, funding acquisition, project administration, supervision, writing – review & editing

## Supplementary materials

**Supplementary Figure 1.**
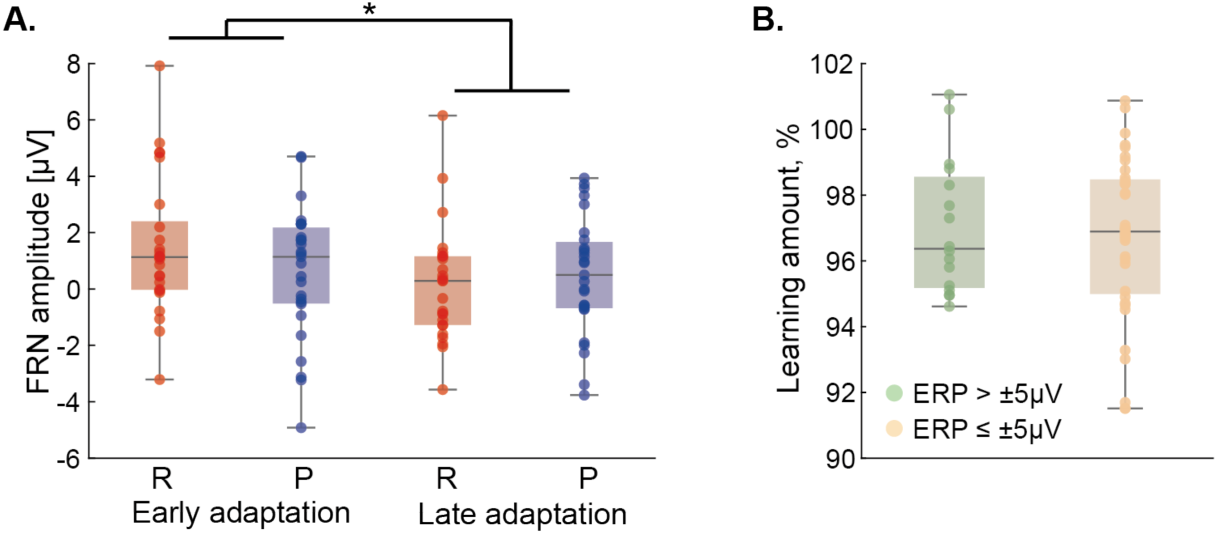
Feedback-related negativity (FRN) results for all participants with valid EEG recordings. (A) FRN amplitudes during the early and late adaptation phases for the reward (R, *n* = 25) and punishment (P, *n* = 27) groups. (B) Comparison of learning amount between participants who exhibited a clear ERP response (greater than ±5 μV, *n* = 16) and those who did not (*n* = 36). In both panels, box plots display the median, interquartile range, and individual data points.

